# Ignoring correlated activity causes a failure of retinal population codes under moonlight conditions

**DOI:** 10.1101/2019.12.18.881201

**Authors:** Kiersten Ruda, Joel Zylberberg, Greg D. Field

## Abstract

The retina encodes visual stimuli across light intensities spanning 10-12 orders of magnitude from starlight to sunlight. To accommodate this enormous range, adaptation alters retinal output, changing both the signal and noise among populations of retinal ganglion cells (RGCs). Here we determine how these light level-dependent changes in signal and noise impact decoding of retinal output. In particular, we consider the importance of accounting for noise correlations among RGCs to optimally read out retinal activity. We find that at moonlight conditions, correlated noise is greater and assuming independent noise severely diminished decoding performance. In fact, assuming independence among a local population of RGCs produced worse decoding than using a single RGC, demonstrating a failure of population codes when correlated noise is substantial and ignored. We generalize these results with a simple model to determine the signal and noise conditions under which this failure of population processing can occur. This work elucidates the circumstances in which accounting for noise correlations is necessary to take advantage of population-level codes and shows that sensory adaptation can strongly impact decoding requirements on downstream brain areas.

## Introduction

Population activity is the currency of sensory systems because individual neurons have limited signal capacity and variable responses to repeated presentations of the same stimuli. This variability is often shared across neurons (termed “noise correlations”), adding a rich complexity to the issue of information processing in neural populations (Averbeck et al., 2006). There is a large body of work showing that these noise correlations can enhance or degrade signaling of sensory information, depending on the structure of noise correlations and their relationship to stimulus-evoked signals (Zohary et al., 1994; Dan et al., 1998; Abbott and Dayan, 1999; Wu et al., 2001; Romo et al., 2003; Zylberberg et al., 2016). A crucial question is how downstream regions can best integrate signals given the noise correlations among their inputs. Perhaps ignoring correlations, or considering input activity as independent, has no adverse effect on computations. On the other hand, downstream regions may need to take correlated activity into account to appropriately process their inputs. Answering this question is critical for understanding how the activity of sensory populations represents stimuli as well as generating informed hypotheses about how downstream circuits process these signals.

In the visual system, populations of retinal ganglion cells (RGCs)—the brain’s sole source of visual information—exhibit activity correlations. Previous work has shown that failing to account for these correlations decreases decoded information by 0-20% (Nirenberg et al., 2001; Pillow et al., 2008; Meytlis et al., 2012). However, these studies were performed under daylight conditions, just part of the retina’s broad operating range that spans 10-12 log units of light intensity. Importantly, the structure of correlated activity changes over light intensities: correlated activity is generally stronger at lower light levels, exhibiting higher peak correlations that extend over longer spatial and temporal scales (Mastronarde, 1983a; Greschner et al., 2011). This shift in correlated activity across populations of RGCs raises the intriguing possibility that light adaptation changes the impact of these correlations on decoding retinal output.

To determine the impact of light adaptation and associated changes in correlated activity, we recorded from populations of rat RGCs with a large-scale multielectrode array (MEA) over conditions spanning rod-mediated (scotopic) to cone-mediated (photopic) light levels. Using a generalized linear model (GLM) to decode retinal activity, we show that at photopic light levels, accounting for correlations among RGCs improves decoding by ~20% compared to assuming the RGCs are independent, similar to previous results in other mammals (Nirenberg et al., 2001; Pillow et al., 2008; Meytlis et al., 2012). However, under scotopic conditions, accounting for correlations showed a significantly larger impact on decoding performance with a ~100% improvement in decoded information. Strikingly, assuming independence across a local population of RGCs produced poorer decoding performance than decoding with a single RGC. In this way, we demonstrate a failure in decoding neural populations when noise correlations are substantial and ignored. Importantly, these results depended on the RGC type that was analyzed, with decoding from OFF-brisk transient RGCs exhibiting greater sensitivity to correlations than decoding from OFF-brisk sustained RGCs. To generalize these results, we created a model of tuned, correlated neurons to identify conditions under which assuming independence causes decoding from the population to perform worse than decoding from a single cell. This model elucidates the circumstances where accounting for correlations not only improves visual processing but is necessary to take advantage of population codes. More generally, this work demonstrates the large impact of context-dependent correlations in sensory processing and raises important questions about how downstream brain areas process retinal signals across light levels.

## Results

### RGCs exhibit greater noise correlations at scotopic light levels

To examine the consequences of pairwise noise correlations on retinal population codes, we recorded RGC responses across a range of light intensities from segments of rat retina on a large-scale MEA (Anishchenko et al., 2010; Ravi et al., 2018). The retina was stimulated with spatiotemporal checkerboard noise to estimate the receptive fields (RFs), contrast response functions and autocorrelation functions of RGCs over the MEA. RGCs were functionally classified according to their light response properties and spiking dynamics, using previously described methods (Yu et al., 2017; Ravi et al., 2018). The results of the classification were validated by observing that each functionally defined RGC type exhibited a mosaic-like organization of RFs that approximately tiled space (Fig. 1A) (Wassle et al., 1981; Devries and Baylor, 1997; Ravi et al., 2018). We initially focus our analysis of correlated activity onto a single cell type: OFF-brisk transient (-bt) RGCs (Fig. 1A). These cells are likely homologous to OFF parasol cells and other transient alpha-like RGCs in other mammals: they exhibit center-surround RFs, short-latency, transient light responses and high contrast sensitivity (Crook et al., 2008; Manookin et al., 2008; Krieger et al., 2017; Ravi et al., 2018). Focusing first on this RGC type facilitated comparing our results to previous work in the primate and rodent retina (Nirenberg et al., 2001; Pillow et al., 2008; Meytlis et al., 2012).

**Figure 1:**
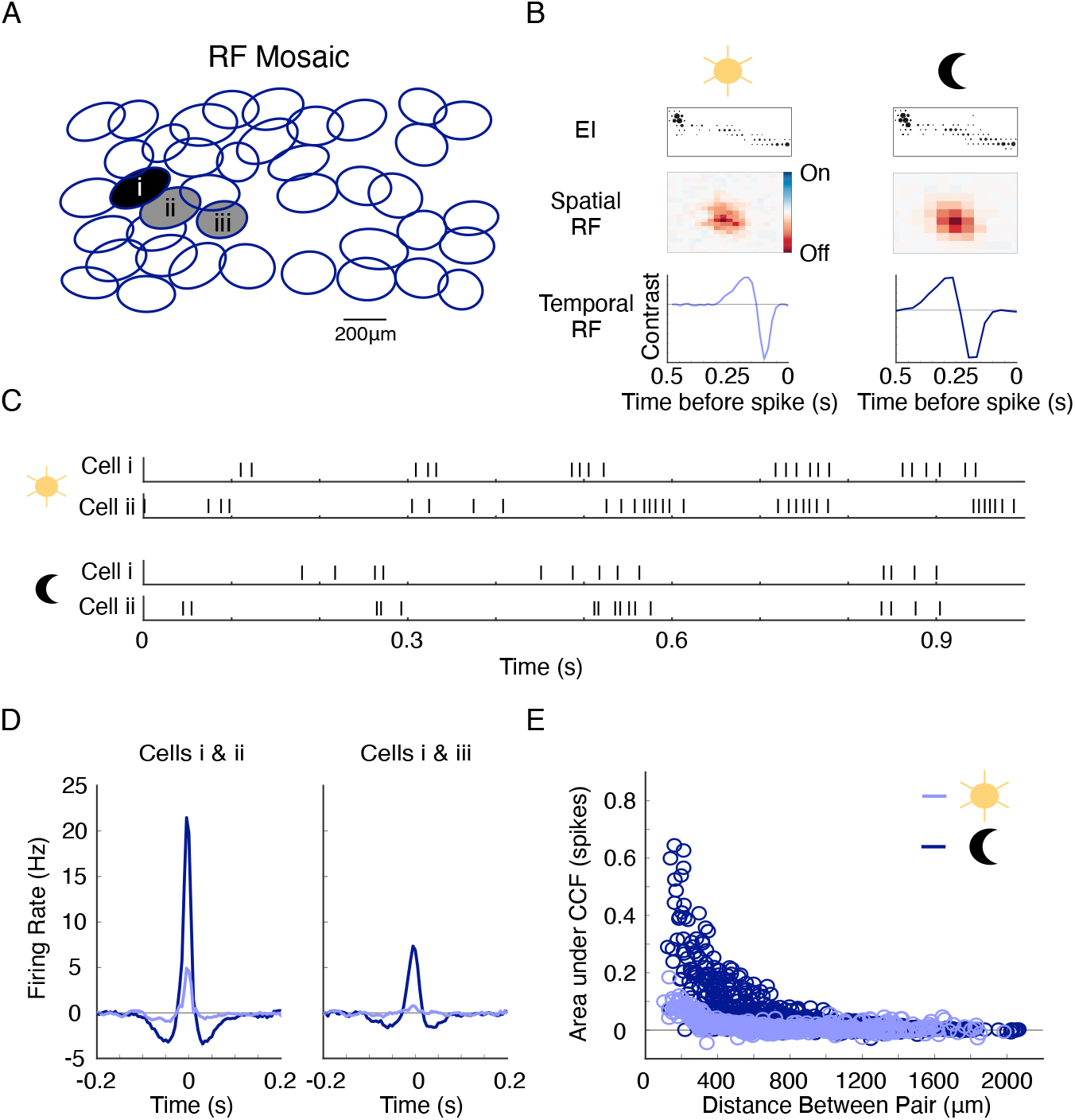
Noise correlation structure and receptive fields of RGCs depend on light level. **A**. Receptive field (RF) mosaic of OFF-bt RGCs. Each ellipse is the 1 SD Gaussian fit to an RGC’s spatial RF. **B**. Top row: Electrical image (EI) of an example cell at two light levels, which enables tracking RGCs across light conditions. Middle row: Spatial RFs at the two light levels. Bottom row: Temporal RF. Both spatial and temporal integration increase in the scotopic condition. **C.** Spike raster of two neighboring RGCs at photopic (top) and scotopic (bottom) light levels responding to white noise stimuli. **D.** Example noise crosscorrelograms across light levels of two primary neighbor cells i and ii in **A** (left) and two secondary neighbor cells i and iii (right). **E.** Strength of noise correlations over pairwise distances for the OFF-bt RGC population. Each point shows the positive area under the crosscorrelation function (CCF) for a given pair of cells. Light adaptation causes expanded correlated noise in time and space for the scotopic condition (561 RGC pairs from 1 retina; see Supp. Fig. 1 for correlated spiking from spontaneous firing).

Understanding the role of light adaptation in retinal coding required tracking the same population of RGCs across rod-mediated (scotopic) and cone-mediated (photopic) conditions. This tracking was achieved by utilizing the electrical image (EI) of each RGC. The EI is computed from the spike-triggered electrical activity of an identified RGC across the MEA (Petrusca et al., 2007). EIs serve as electrical footprints of each cell and are stable despite changes in responses across light levels (Field et al., 2009) (Fig. 1B). This tracking procedure was further validated by observing a nearly identical mosaic-like organization of RFs across the scotopic (1.0 Rh*/rod/s) and photopic (10,000 Rh*/rod/s) light levels examined in these experiments (see Methods).

The pairwise noise correlations among OFF-bt RGCs were greater under the scotopic condition (Fig. 1D & E). We computed all cross-correlograms between OFF-bt RGCs responding to the white noise stimulus and estimated noise correlations by removing stimulus-induced correlations (see Methods). The area under the peak and width of the noise correlations between primary neighbors were greater under the scotopic conditions (Fig. 1D; Table 1). The spatial scale of correlations over the population of OFF-bt RGCs was also larger at the scotopic light level (Fig. 1E; Table 1). To verify that these noise correlations are not critically influenced by the white noise stimulus, we also considered correlated noise during spontaneous activity (Supp. Fig. 1). Those measurements revealed similar changes in correlation structure across light levels. Cumulatively, these observations indicate higher magnitude correlations that have broader temporal and spatial scales across the population of OFF-bt RGCs at the scotopic light level, consistent with previous studies (Mastronarde, 1983a; DeVries, 1999; Greschner et al., 2011). In the subsequent sections we utilize a model-based decoding approach to determine the impact these changes in correlation structure have on decoding visual stimuli from populations of OFF-bt RGCs.

**Table 1:** Measurements of correlation structure across light levels for the two RGC types. Values are mean ± s.e.m. All data comes from 4 retinas for the photopic condition and 3 retinas for the scotopic condition (3 retinas in common between conditions).

| | | CCF area (spikes) | CCF width (s) | Correlation spatial scale (μm) | n (primary RGC pairs) |
| --- | --- | --- | --- | --- | --- |
| OFF-bt | Photopic | $0.105 \pm 0.004$ | $0.037 \pm 0.001$ | $205 \pm 4$ | 128 |
| | Scotopic | $0.253 \pm 0.013$ | $0.045 \pm 0.001$ | $280 \pm 4$ | 99 |
| | Change over light levels | $0.165 \pm 0.001$ | $0.012 \pm .001$ | $74 \pm 5$ | 99 |
| OFF-bs | Photopic | $0.029 \pm 0.003$ | $0.033 \pm 0.001$ | $141 \pm 6$ | 96 |
| | Scotopic | $0.063 \pm 0.005$ | $0.035 \pm 0.001$ | $167 \pm 5$ | 72 |
| | Change over light levels | $0.037 \pm 0.001$ | $0.013 \pm 0.002$ | $26 \pm 8$ | 72 |
| P values |  |  |  |  |  |
| OFF-bt: photopic vs scotopic | | $p \ll 0.001$ | $p \ll 0.001$ | $p < 0.005$ | |
| OFF-bs: photopic vs scotopic | | $p \ll 0.001$ | $p = 0.29$ | $p < 0.05$ | |
| Photopic: OFF-bt vs OFF-bs | | $p \ll 0.001$ | $p < 0.01$ | $p < 0.001$ | |
| Scotopic: OFF-bt vs OFF-bs | | $p \ll 0.001$ | $p \ll 0.001$ | $p < 0.001$ | |
| Light level change: OFF-bt vs OFF-bs | | $p \ll 0.001$ | $p = 0.63$ | $p < 0.005$ | |

### RGC responses are fit well by the GLM across light levels

The model-based decoding approach we used involves first fitting an encoding model to capture the relationship between visual stimuli and RGC spiking. This model will be inverted to estimate stimuli given RGC spike trains. Importantly, we are not claiming that this exact model-inversion procedure is used in brain areas downstream of the RGCs. Since the exact computations downstream of the retina are unknown, we chose an optimal decoding approach. This procedure yields a way to estimate how well an ideal downstream system could estimate the stimulus, given the RGC spike trains using different assumptions about correlations between cells (Averbeck et al., 2006; Pillow et al., 2008).

To quantitatively describe RGC spiking in response to a checkerboard stimulus, we use the Generalized Linear Model (GLM), a phenomenological model for retinal encoding that can also be used for Bayesian decoding (Pillow et al., 2008). The GLM transforms visual stimuli to spike times by first filtering the stimulus through the spatiotemporal RF and applying a spike history filter to account for refractoriness and spike bursts (Fig. 2A). This signal is then passed through a static nonlinearity to yield a predicted firing rate, and spike times are generated with a Poisson process. We first fit OFF-bt RGCs with an independent version of the GLM, in which each cell is fit individually and the spiking of one RGC is independent of the other RGCs (except for stimulus-induced correlations). Cells were fit at each light condition separately to optimize model performance at each light level. The independent GLM predicted held-out responses well at both light levels, as measured by the explained variance in firing rates (photopic: 0.59 ± 0.01, mean ± s.e.m., 97 cells from 4 retinas, scotopic: 0.58 ± 0.01, 66 cells from 3 retinas; Fig. 2C, D). Furthermore, the GLM captured changes known to occur in light adaptation, such as larger spatial RFs, slower temporal integration, and expanded interspike interval distributions at the scotopic light level (Barlow et al., 1957; Enroth-Cugell and Shapley, 1973) (Fig. 1B and Supp. Fig. 2).

**Figure 2:**
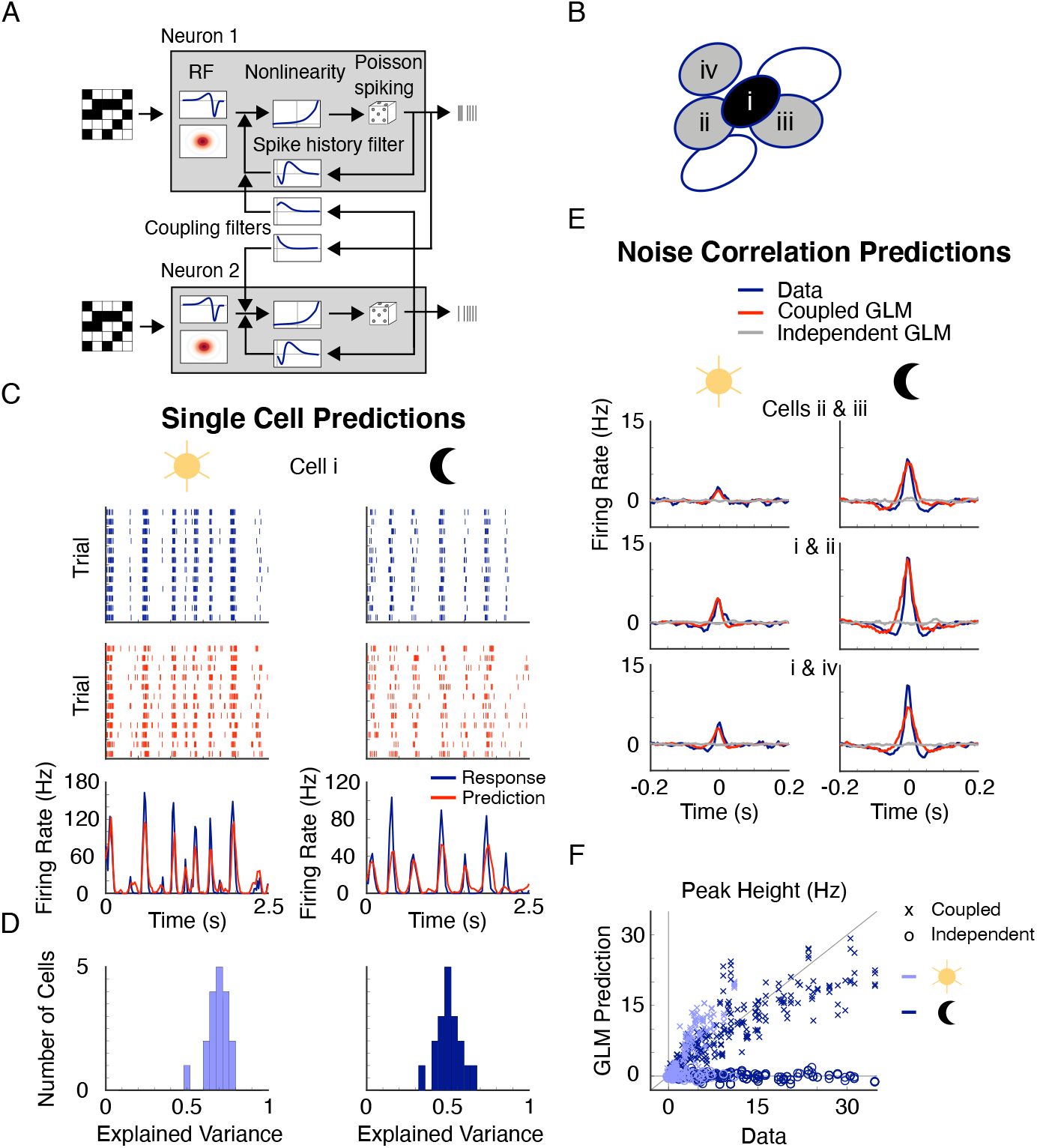
A generalized linear model (GLM) captures RGC responses and pairwise correlation structure. **A.** GLM diagram for two coupled cells, adapted from Pillow et al. 2008. The stimulus is filtered by a RF, this generator signal is passed through a nonlinearity to generate firing rates, and then stochastic spikes are created. Spike history and coupling filters (for the coupled GLM) also influence the generator signal. **B.** Example local group of RGCs to which the GLM model is fit. **C.** Example recorded raster (blue), GLM predicted raster (red), and PSTH and predicted PSTH (bottom) for cell i in **B**. Left column shows photopic light level and right column shows scotopic light level. **D.** Distribution of explained variances for the GLM predicted PSTHs for photopic (left) and scotopic (right) light levels (18 Off-bt RGCs from 1 retina). **E.** CCFs and GLM predicted CCFs of cell pairs shown in **B** at two light levels. **F.** The coupled GLM predicts close to the measured noise correlation values, while the independent GLM predicts no noise correlations (189 pairs from 11 groups of RGCs from 1 retina; see Supp. Fig. 2 legend for data from all retinas).

To account for correlations between RGCs and determine their impact on decoding, we separately fit a coupled version of the GLM. The coupled GLM includes pairwise coupling filters so that the activity of one RGC can influence the responses of other RGCs, allowing the coupled GLM to capture noise correlations in RGC activity (Pillow et al., 2008). Because correlations decrease rapidly with distance between pairs of cells (Fig 1E), we used local groups of RGCs in the coupled GLM, choosing each group based on a central RGC and all of its recorded neighbors (Fig. 2B). For single cell PSTHs, the coupled GLM predictions and performances were very similar to those of the independent GLM at both light levels (Supp. Fig. 2). The coupled model predicted noise correlations well, while the independent model did not predict any, as expected (Fig. 2E, F). Having established the GLM as an accurate description of RGC activity under scotopic and photopic conditions, we next use the independent and coupled versions to probe the impact of correlations on decoding retinal output over light levels.

### Scotopic decoding performance is severely decreased when RGC correlations are ignored

We estimated white noise stimuli from recorded responses to elucidate the impact of correlations on processing OFF-bt RGC output. To perform model-based decoding of responses, we inverted the independent and coupled GLMs fit to recorded OFF-bt RGCs (see Fig. 3). We compared the decoding performance between these two models to determine the extent to which ignoring noise correlations between RGCs diminished decoding performance. We performed Bayesian decoding, which optimally extracts stimulus information available in the RGC response structure that is captured by the GLM (Pillow et al., 2008). Given a set of spike times from a local group of RGCs, we decoded the intensity of a single stimulus pixel over six sequential frames. For this analysis, the stimulus pixels and RGCs were chosen such that the pixel was predominantly covered by the center-most RGC of the group of cells (see Methods). We report decoding performance with a signal-to-noise ratio (SNR), which quantifies the information rate in bits/s that the decoded estimate provides about the actual stimulus (Warland et al., 1997).

**Figure 3:**
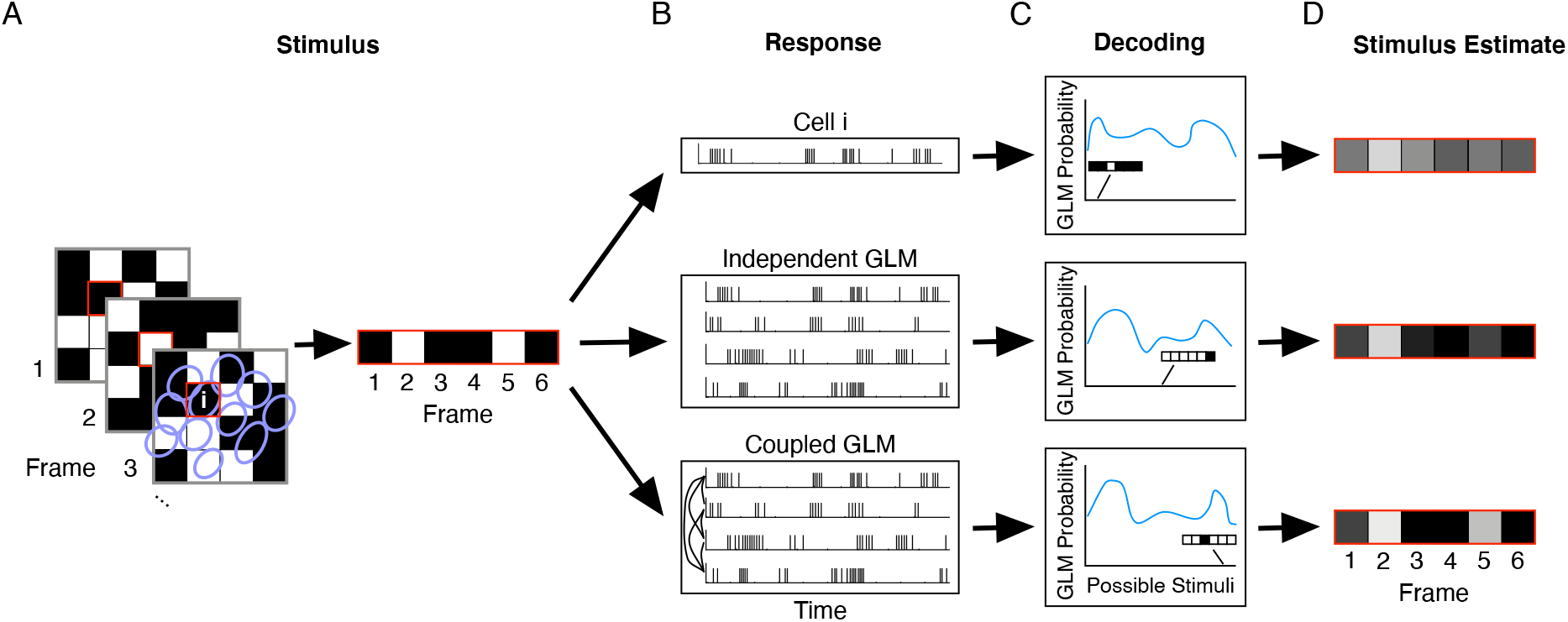
Schematic showing GLM-based Bayesian decoding. **A.** One stimulus pixel, highlighted in red, is chosen over several frames (left), yielding a sequence of intensity values (right). **B.** The corresponding response of a single RGC (top) or population of RGCs (middle and bottom) are extracted. **C.** The probability of each possible stimulus given the input response is computed under a GLM fit to that population. **D.** Summing over the possible stimuli weighted by their probabilities gives the optimal Bayesian estimate of the stimulus. In general, each GLM provides a different estimate of the original stimulus.

At the photopic light level, the coupled GLM is a more accurate decoder, providing 22 ± 3 % (mean ± s.e.m.) more information than the independent GLM over all groups of OFF-bt RGCs (Fig. 4A, B; 55 groups of RGCs from 4 retinas). However, at the scotopic light level, the importance of correlations for accurate decoding substantially increased for OFF-bt RGCs. Accounting for noise correlations with the coupled GLM provided 104 ± 18 % more information than the independent GLM (Fig. 4B; 37 groups of RGCs from 3 retinas, difference over light levels p << 0.001). Furthermore, the improvement in decoding for a given group of cells correlated positively with the strength of noise correlations in that group, indicating that accounting for correlated activity enhances decoding most when correlated noise is largest (Fig. 4C).

**Figure 4:**
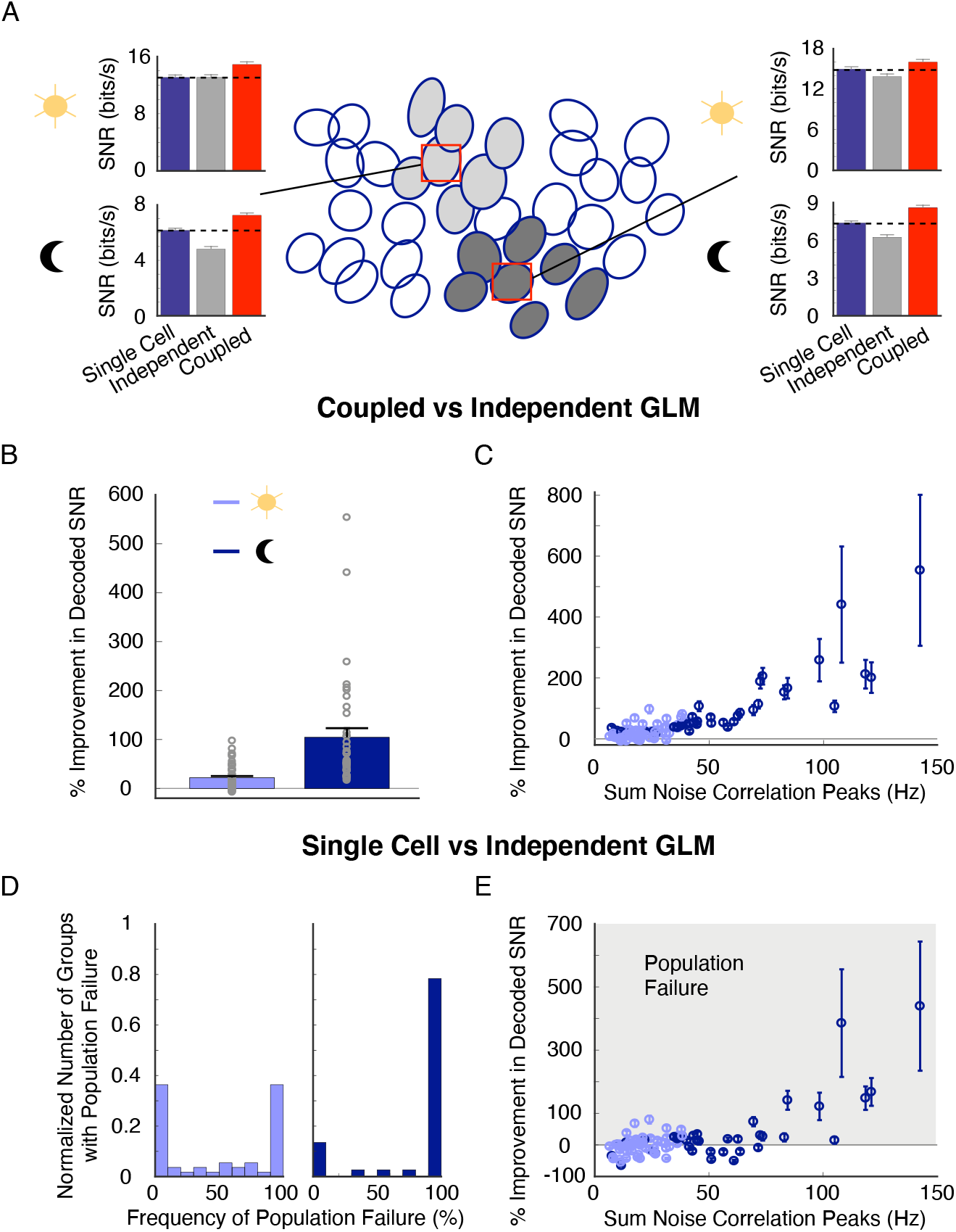
Assuming population independence yields poor decoding performance under scotopic light levels and can perform worse than decoding individual RGC responses. **A.** Decoding examples for two groups of OFF-bt RGCs. In these examples the coupled and single cell GLMs decode better than the independent GLM at scotopic light levels: see dashed lines in bar plots. Error bars are from bootstrapping decoded SNR. **B.** Average percent improvement in decoded SNR at each light level when using the coupled GLM over the independent GLM. **C.** Percent improvement in decoding relates positively to the amount of noise correlation in the groups of RGCs (R^2^ = 0.6 for a single term exponential). Noise correlation for each group of RGCs is quantified by accumulating the peaks of CCFs between the centered RGC and its neighboring cells. **D.** Distribution of frequencies of the single cell GLM decoding better than the independent GLM, termed population failure. Frequency of population failure for each group is computed over bootstraps. **E.** Population failure relates positively to the amount of noise correlation. For panels **B-E**: photopic: 55 groups of RGCs from 4 retinas, scotopic: 37 groups of RGCs from 3 retinas (3 retinas in common between conditions).

### Population failure: Single RGCs can outperform populations when assuming independence

To better understand the significance of the information loss due to ignoring correlations, we compared the decoding performance of the independent GLM to that of the best-performing single cell model. This single cell model was simply the individual GLM for the RGC centered over the decoded pixel. Surprisingly, in many of the tested groups, the single cell GLM outperformed the independent population GLM (Fig. 4A). We call this effect ‘population failure’ because the GLM fit to a population of RGCs decodes less information than from a single RGC when the population is assumed to be independent.

Population failure primarily occurred at the scotopic light level (Fig. 4D). In that condition, the majority of groups of RGCs exhibit this population failure mode (83 ± 5.7 %, mean frequency of population failure ± s.e.m., 37 groups of RGCs, photopic: 50 ± 5.9 %, 55 groups of RGCs), and among those groups the single cell GLM provided 73 ± 23 % more information than the independent GLM (Fig. 4E; photopic: 19 ± 4 %). Notably, the single cell GLM uses the exact same parameters as its corresponding cell in the independent GLM, so our findings are not a consequence of model fitting issues. Rather, this result demonstrates that decoding under the assumption that a population of RGCs is independent can be so suboptimal that it extracts less information than a single cell. This population failure under the assumption of independence is a striking example of the importance of accurately accounting for correlations in processing population activity, particularly in scotopic conditions.

We next performed a series of controls to assess how particular details of our decoding analysis might influence these results. In the analyses above, we chose local groups of cells based on a central RGC with its nearest neighbors. There the majority of the RFs over the population of RGCs had some overlap with the decoded stimulus pixel so that each cell provided nonzero decoding information about that pixel intensity (e.g. Fig 4A; note RF outlines are plotted at a 1 SD contour of a Gaussian fit, so the RFs extend well beyond the RF outline). To determine how this choice of population impacts decoding, we also decoded using larger groups of cell clusters, including secondary and tertiary neighbors. Including RGCs with RFs far away from the decoded stimulus pixel did not significantly alter the performances of the coupled or independent GLMs because those cells contribute minimal information to decoding and do not exhibit strong correlations with RGCs close to the decoded stimulus pixel, as expected (Supp. Fig. 3). Thus, our selection of local groups of RGCs is not a crucial factor in the role of correlations for decoding.

We further sought to ascertain whether population failure generalizes beyond temporal decoding by instead decoding spatial patterns of stimulus pixels for one movie frame. Under this decoding task, the coupled GLM continues to perform substantially better than the independent GLM at the scotopic light level (52 ± 16 %, mean ± SD over bootstraps; Supp. Fig. 4). In addition, the independent GLM decodes less information than smaller groups of coupled RGCs, exhibiting population failure because 18 cells in the independent GLM perform worse than 7 cells in a coupled GLM. These results demonstrate that the large cost of ignoring correlations is a general feature of spatial and temporal decoding from OFF-bt RGCs.

Finally, to verify that changes in correlation structure causally affect the difference in decoding performance between coupled and independent GLMs, we simulated RGC population responses with the GLM and then used the GLM to decode these simulated responses. As we observed when decoding measured responses at the scotopic light level, we hypothesized that stronger coupling among neurons would lead to a higher percent improvement in decoding SNR when accounting for noise correlations versus assuming independence. Indeed, larger correlation strength between RGCs causes the coupled decoder to perform much better than the independent decoder (Supp. Fig. 5). This simulation emphasizes how the amount of correlated noise impacts decoding performance under the independence assumption.

### The cost of ignoring correlations depends on RGC type

We next investigated the extent to which the population failure phenomenon occurs in a distinct RGC type, the OFF-brisk sustained (-bs) RGCs. These cells likely correspond to RGCs called OFF delta or OFF sustained alpha cells in other studies (Manookin et al., 2008; Krieger et al., 2017; Ravi et al., 2018). The correlation structure across the OFF-bs RGC population shows that the magnitude, timescale and spatial scale of correlations is smaller than in OFF-bt RGCs (Fig. 5A; Table 1). In addition, the correlations among OFF-bs RGCs do not change with light adaptation as much as in OFF-bt RGCs (Table 1). To determine the role of accounting for correlations in decoding OFF-bs RGC activity, we next compared independent and coupled GLM decoders fit to groups of OFF-bs RGCs (Fig. 5). Accounting for correlations only improved decoded SNR by 2.9 ± 0.7 % in the photopic condition and 4.4 ± 0.8 % in the scotopic condition (Fig. 5C; photopic: 37 groups of RGCs from 4 retinas, scotopic: 20 groups of RGCS from 3 retinas, difference over light levels p = 0.45). While some single cell GLMs decode better than the independent GLM, the frequency and amount of this population failure under the independence assumption was much smaller than in OFF-bt RGCs (Fig. 5E, F; photopic: frequency of population failure = 19 ± 5 %, mean ± s.e.m., % improvement when there is population failure = 1.9 ± 1 %, scotopic: frequency of population failure = 28 ± 8 %, % improvement when there is population failure = 0.7 ± 0.2 %). These results demonstrate that the role of noise correlations in decoding RGC activity depends on both adaptation state and cell type.

**Figure 5:**
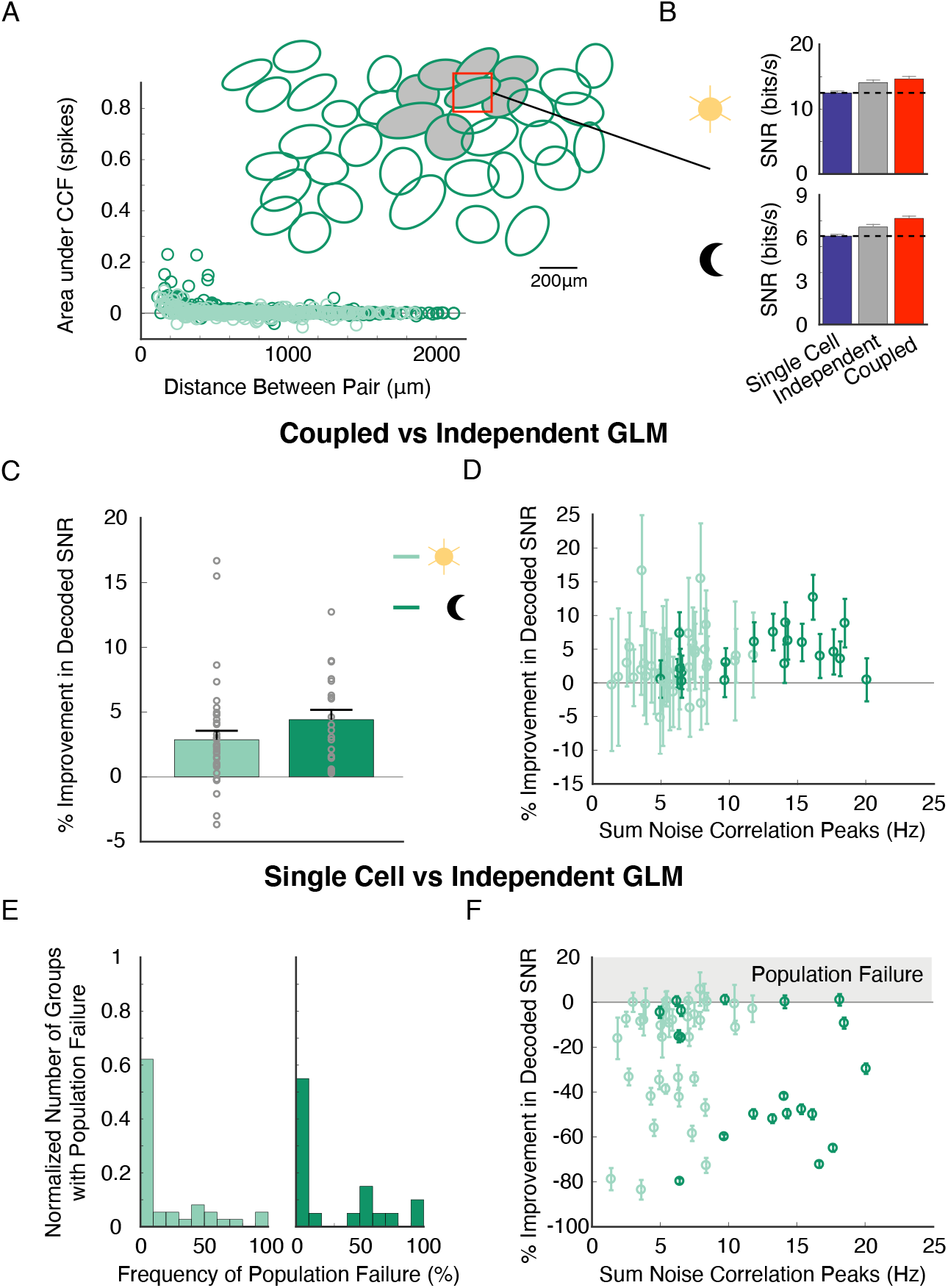
Diminished decoding performance when assuming independence depends on RGC type. **A.** Correlation structure of OFF-bs RGC population across light levels (left) and RF mosaic (right) in the same retina as Fig. 1 (351 RGC pairs). **B.** Decoding result for an example group of cells over light levels. In this instance the independent and coupled GLMs perform similarly. **C.** Average percent improvement in decoded SNR across light levels when using the coupled GLM over the independent GLM. Note the compressed y-axis compared to Fig. 4B. **D.** Relationship between percent improvement in decoding and summed noise correlations between a centered RGC and its neighboring cells. Note the compressed x-axis compare to Fig. 4C. **E.** Distribution of population failure frequencies. **F.** Population failure as a function of the amount of noise correlation in a group of RGCs. For panels **C-F**: photopic: 37 groups of RGCs from 4 retinas, scotopic: 20 groups of RGCs from 3 retinas (3 retinas in common between conditions).

### A simple geometric model reproduces population failure

Under the scotopic condition, assuming noise independence among OFF-bt RGCs frequently caused population failure. To provide an intuition for this potentially counterintuitive result, we utilized a previously developed geometric visualization of noise correlations and decoding performance (Averbeck et al., 2006) (Fig 6). We created a simplified model of two neurons responding to two stimuli, using the d prime metric to quantify how well the neurons could discriminate the stimuli with their firing rates (Averbeck and Lee, 2006). In one case, the two cells exhibit strong noise correlations causing elliptical joint response distributions (Fig 6A, green and blue solid ellipses). The optimal decoder (red line) accurately discriminates the two populations (Fig 6C, red bar). However, if the noise is assumed to be independent between the two cells (Fig 6A, green and blue dashed circles), the decoder is nearly orthogonal to the optimal decoder (compare black and red lines). This independence assumption causes a large decrease in decoding performance (Fig 6C, light blue bar). Next, we discriminated the two stimuli using just the response distributions for cell 1 (Fig 6B). In this example, the single cell outperforms the two-cell decoder that assumes independence among the cells (Fig 6C). In a second case, the noise correlations are weaker, resulting in a close similarity between the decoder under the independence assumption and the optimal decoder (Fig 6D, black and red lines). Assuming independence causes a very small decrease in discrimination performance and outperforms discrimination from a single cell (Fig 6E, F). This model demonstrates how population failure can occur in a very simple system—where decoding from a single neuron performs better than decoding from two neurons with non-zero stimulus information. Between these two cases we examined (Fig 6A, D), we only changed the strength of the noise correlations and held constant all other parameters, namely, the stimulus-dependent firing rates and variances of the two neurons. Thus, changes in noise correlations alone can cause conditions under which population failure occurs. In general, however, the relative sign and strength of the signal and noise correlations are the key elements in determining the consequences of decoding correlated responses under the independence assumption (Averbeck et al., 2006). In the next section we examine how these parameters shape decoding performance in a population model that more closely relates to our experiments.

**Figure 6:**
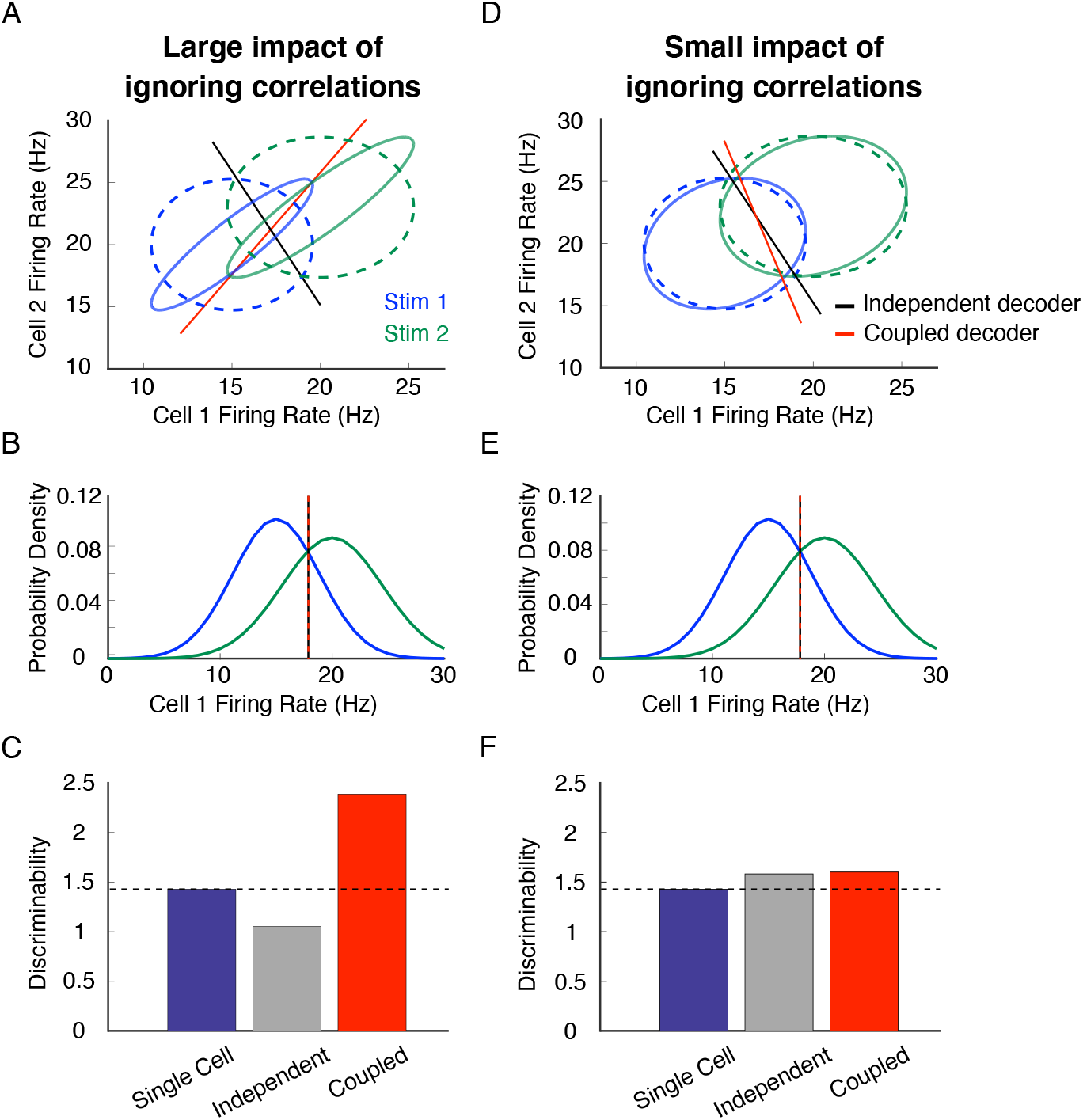
A simple, two-cell model can reproduce population failure when assuming independence in a correlated population. **A.** Joint responses when correlated activity is taken into account (solid ellipses) and when it is ignored (dashed ellipses). Geometrical perspective on decoding adapted from Averbeck et al. 2006. The optimal decoder with knowledge of correlations (red line) is very different than the decoder without knowledge of correlations (black line), resulting in high error for the independent decoder. **B.** Response distributions of cell 1 in panel **A**, which reveal the optimal decoding line for a single cell model lies at x = 18 Hz. **C.** The independent decoder (**A**, black line) discriminates the two stimuli worse than the coupled decoder (**A**, red line) and the single cell decoder (cell 1, **B**). **D-F.** same as **A-C** but with low noise correlations. Here the independent decoder is similar to the optimal decoder, resulting in similar discrimination performance.

### Receptive field overlap, correlation strength and its spatial scale dictate population failure

To investigate the conditions under which a single RGC can outperform a population that is assumed to be independent, we modeled our experimental findings by simulating a two-dimensional grid of RGCs (Fig 7A). We systematically varied three parameters that determine correlation structure across this population: peak correlation strength, the spatial scale of correlation, and RF overlap (Fig. 7A). The model consists of linear RGCs responding to a white noise stimulus. We arranged the RFs in a hexagonal grid to approximate the mosaic of one RGC type, with the relationship between RF and stimulus pixel sizes set similarly to those in our experiments. We again used discriminability (d prime) to quantify how well intensity values in the central stimulus pixel can be discriminated given the neural responses (Averbeck and Lee, 2006). As with our GLM-based approach, we compared the performance of a decoder that accounts for correlations among cells (coupled decoder), one that assumes independence among cells (independent decoder), and one that just uses the responses of one RGC (single cell decoder).

**Figure 7:**
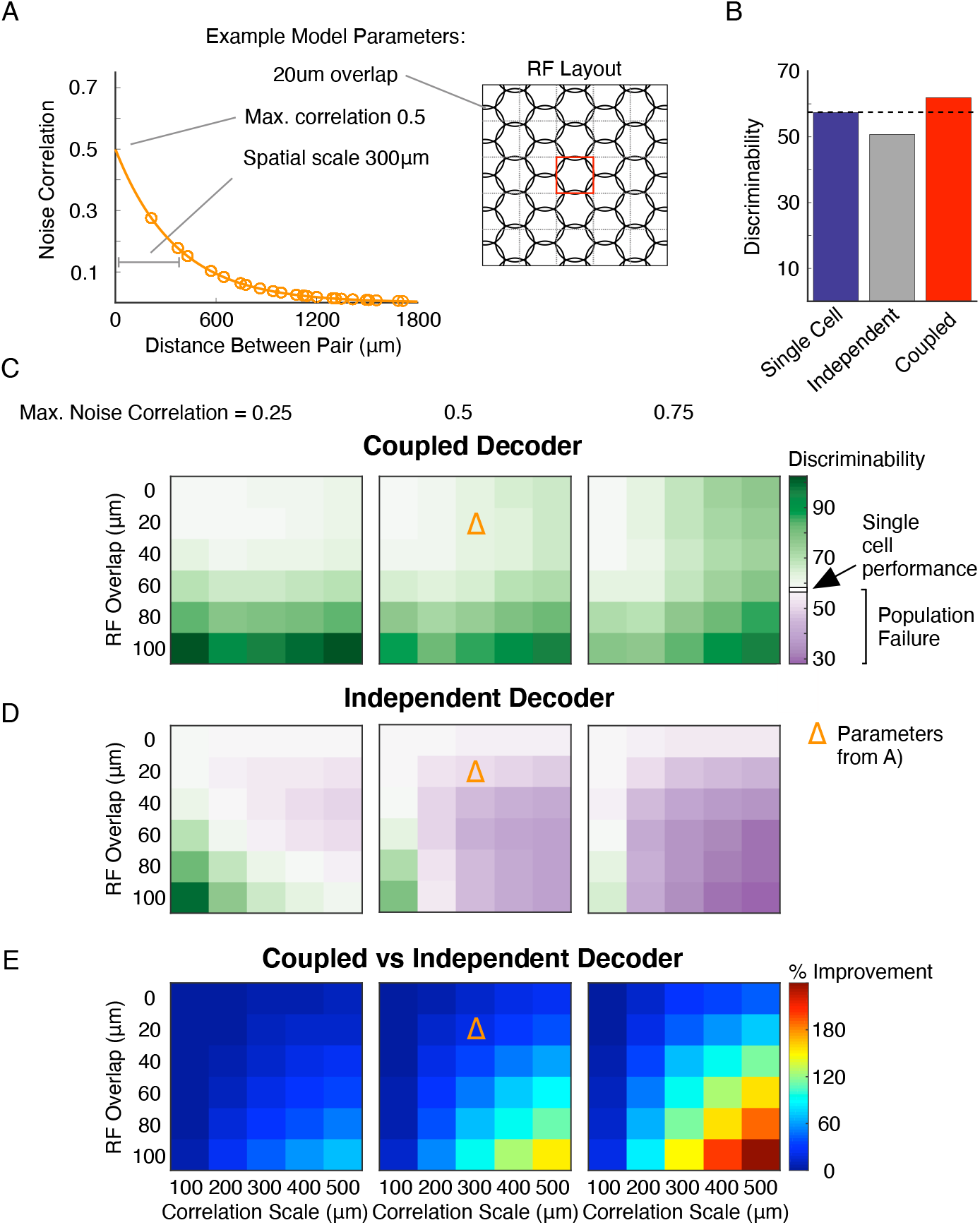
RF overlap, peak correlation, and spatial extent of correlations dictate the conditions under which accounting for correlations is necessary for population failure. **A.** Correlation structure (left) and RF mosaic (right) of an example model with the given RF overlap, maximum correlation, and spatial scale of correlations. **B.** Decoding results from the model parameters shown in **A** illustrate a state in which the assumption of independence causes the population to perform worse than a single cell. **C.** Color map of discriminability for the coupled decoder over all values for RF overlap, maximum noise correlation, and spatial scale of noise correlations. **D.** Same as **C** but for the independent decoder. As the magnitude and scale of noise correlations increases, the independent decoder discriminates less information. Purple regions indicate states in which the population decoding assuming independence performs worse than decoding from a single cell. **E.** Percent improvement in discriminability achieved by the coupled decoder over the independent decoder.

When there is little overlap between RFs and noise correlations are present, the independent decoder often discriminates the stimulus worse than the single cell model (Fig. 7D, top rows, purple areas). The cost of assuming independence in the population becomes more severe as the noise correlations are made stronger and/or broader. As RF overlap increases, the independent decoder performs better than the single cell decoder when correlations have a small spatial scale and magnitude (Fig. 7D, left columns). This improved performance results from neighboring cells providing more signal about the intensity of the decoded pixel, which overcomes the errors due to ignoring small noise correlations. However, ignoring larger and broader noise correlations eventually outweighs this advantage, resulting in more extreme population failure (up to 50% less discriminability than the single cell decoder for the parameters we explored; Fig. 7D, right columns). Note that the coupled decoder discriminates much better than the independent decoder in the presence of strong and broad noise correlations (Fig. 7E). This simplified model highlights that accounting for correlated noise is most important for decoding the stimulus when correlations are large, in agreement with our experimental findings. By considering the transition into population failure modes based on noise correlation parameters, this model demonstrates how changing correlation structure in OFF-bt RGCs across light levels can alter the frequency and magnitude of population failure. The correlations in OFF-bs RGCs, however, are generally too small at both light levels for population failure to occur.

These modeling results also reproduce a point about the limits that noise correlations can place on a neural system. Focusing only on the correlated decoder, there are several conditions where high noise correlations limit the discriminability of the neural population compared to weak noise correlations (Fig. 7C, bottom rows). This phenomenon has been previously described, and since we are primarily focused on the consequences of assuming independence given the presence of noise correlations, we do not consider it further (Averbeck et al., 2006).

## Discussion

A major question in early vision is how circuits downstream of the retina process the visual information conveyed by populations of RGCs. Central to this question is the impact of correlated activity among RGCs, which can be a significant factor in neural computations depending on context. Here we examine how light adaptation alters the role of correlations in decoding visual stimuli from RGC populations in the rat retina. We find that under moonlight conditions, decoders assuming independent responses among OFF-bt RGCs recover much less visual information than decoders that account for pairwise correlations. This reduction in performance can be so large that decoders assuming independence perform worse than decoding from a single RGC (Figs 4, 6, and 7). We call this state ‘population failure’ because decoding the population fails to reach the performance of a single cell. Accounting for correlations, however, avoids this state and enables decoders to benefit from population codes. We use a simple model to demonstrate how the structure of activity correlations determines the cost of assuming independent responses, accounting for why our results depend both on light level and RGC type. These findings raise several questions about the role of correlations in adaptation and visual processing that we discuss below.

### Comparison to previous studies, interpretation and caveats

The importance of correlated activity is a much-debated topic in vision research (Dan et al., 1998; Nirenberg et al., 2001; Schneidman et al., 2003; Schnitzer and Meister, 2003; Puchalla et al., 2005; Montani et al., 2007; Pillow et al., 2008; Graf et al., 2011; Berens et al., 2012; Meytlis et al., 2012). Previous studies have examined the role of correlated spiking in both visual encoding and decoding, yielding a range of conclusions. For decoding, studies have concluded that between 0-40% more information is available when decoders account for correlations (Dan et al., 1998; Nirenberg et al., 2001; Pillow et al., 2008; Meytlis et al., 2012). Our results are most comparable to the Pillow et al. 2008 and Meytlis et al. 2012 studies because they analyzed similar population sizes and employed the same GLM-based decoding strategy. The decoding improvement we find at the photopic light level agrees relatively well with their 20% and 13% results, respectively. Our study departs from previous work by determining how this decoding improvement depends on adaptation state and cell types that encode distinct visual features (OFF-bt vs OFF-bs RGCs). The effect of light level on OFF-bt RGCs is particularly striking: decoded information can be doubled by accounting for correlations. This improvement is a substantially larger effect than previous results at photopic light levels, illustrating the potent impact of light adaptation on retinal output.

How could accounting for correlations improve retinal decoding? One possibility is that correlated activity conveys visual features that are unavailable from individual responses, such as fine spatial features at the intersection between two RFs (Meister et al., 1995; Meister, 1996; Dan et al., 1998). To check for this possibility, we analyzed synchronous spike triggered averages (sSTAs) from pairs of RGCs. We did not find evidence that synchronous spikes provide a higher acuity representation of visual space (Supp. Fig. 6). An alternative possibility is that accurate decoding requires an accurate model of the noise in RGC populations (Averbeck et al., 2006). When correlated noise is large and spatially extensive, such as for OFF-bt RGCs at scotopic light levels, assuming independence is the wrong noise model, and this assumption diminishes decoding so much that performance can fall below that of decoding from a single cell.

A simple intuition for the population failure effect can be achieved by considering the following situation. If a single OFF-bt RGC generates a brief volley of spikes, a decoder will interpret this response as resulting from a transient decrease in light intensity. If all the OFF-bt RGCs around that cell also generated spikes, the decoder will estimate a large decrease in light intensity because many cells were driven to spike together. However, this interpretation may only be correct if the OFF-bt RGCs are acting independently. If the decoder knows the cells are strongly correlated, then it should discount this conclusion in favor of a smaller decrease in light intensity.

Many of the findings presented here are based on GLM fits to the responses of RGC populations, raising the possibility that at least some of these conclusions are model-dependent. The GLM captures a majority of the response variance to white noise stimuli (up to 80%), but remains an imperfect model of RGC encoding (McFarland et al., 2013; Heitman et al., 2016). This mismatch between model and data is likely to impact the quantitative estimates that we and others have made on the cost of assuming RGCs are independent (Pillow et al., 2008; Meytlis et al., 2012). Of particular concern is whether assuming independence in a population of RGCs can actually yield worse performance than decoding a single RGC. To address this issue, we also utilized a more general examination of how correlations can impact decoding. Fig. 6 demonstrates that population failure is certainly possible. Fig. 7 shows that this effect depends on the amount of RF overlap and strength of noise correlations, both of which change with light level. This simplified model has many differences from our data, with uniform, circular RFs, uniform RF overlap, firing rates that depend linearly on the stimulus, and decoding using discriminability (d prime) rather than GLM-based optimal stimulus estimation. Nevertheless, the simplified model in Fig. 7 reproduced the trends in our data. Furthermore, we show that population failure can occur when decoding spatial stimulus patterns from RGC responses (Supp. Fig 4), indicating that these results are not specific to temporal decoding. Together, these analyses show that ignoring strong correlations can degrade decoding and reproduce population failure in a manner that does not depend strongly on the details of the decoding task or the precise nature of the RGC output.

### Light adaptation

Light adaptation crucially influences how retinal circuits encode visual scenes. Between scotopic and photopic light levels, input to RGCs switches from rod-to cone-mediated pathways. This circuit switch alters both single RGC response properties and correlated activity. For individual RGCs, spatial and temporal integration increases under scotopic conditions (Barlow et al., 1957; Enroth-Cugell and Shapley, 1973). Other aspects of RGC activity also depend on light level, including the polarity of stimuli that drive responses, firing rates, and the extent to which spatial integration is linear (Barlow and Levick, 1969; Grimes et al., 2014; Tikidji-Hamburyan et al., 2015). The switch from rod-to cone-mediated circuits also results in altered common input to RGCs, one of the underlying causes of RGC correlations (Mastronarde, 1983a, b; Greschner et al., 2011). In general, weaker RF surrounds in scotopic conditions result in greater overlap between RF centers and thus more common input between neighboring RGCs. Furthermore, at the low light level used here (1R*/rod/s), AII amacrine cells are expected to be extensively coupled by gap junctions (Bloomfield and Volgyi, 2004), which would also tend to increase the amount of common input between nearby RGCs. Finally, a subset of RGC types are electrically coupled (Völgyi et al., 2009), and the strength of this coupling can be altered by light level (Hu et al., 2010). Thus, there are several mechanisms by which light adaptation can strongly impact correlated spiking.

These changes in signal and noise across light levels raise the question of how light adaptation influences information across populations of RGCs. Efficient coding theory—the idea that sensory systems are optimized to encode natural stimuli—has been successful at explaining why RF structure changes across light levels (Attneave, 1954; Barlow, 1961; Atick and Redlich, 1990; Van Hateren, 1993). However, this theory assumes that RGCs do not exhibit correlated noise, much less that this correlated noise changes with light level. Therefore, a useful direction for future examinations of efficient coding theory is to determine how light-level dependent changes in correlated activity impact predictions about the optimality of adaptation in RGC responses.

Our results showing that the importance of noise correlations in decoding can change across light levels highlight how the role of correlated activity in neural computations can be altered by context. This contextdependent change parallels work demonstrating modulation of correlations in other brain regions. For example, attention has been shown to alter the structure of noise correlations in cortical areas such as MT and V4, resulting in more informative population activity and likely causing improved behavioral performance on visual discrimination tasks (Cohen and Maunsell, 2009; Mitchell et al., 2009). Note that unlike the present work, these studies assumed a downstream computation that averages population activity, yielding increased information when noise correlations are reduced because correlated noise cannot be averaged away (Zohary et al., 1994). However, the general point still stands that context can change the structure of signal and noise correlations, which in turn impacts circuit computations.

### Implications for downstream processing

Our results highlight several implications for how downstream circuits may process retinal output. Although the impact of ignoring RGC correlations may depend on particular post-retinal computations, assuming independence among correlated RGCs likely reduces information that can be extracted from retinal activity. Even in the context of a simple linear readout of RGC responses, ignoring strong noise correlations among RGCs can result in suboptimal weighting of RGC inputs compared to weighting determined by an accurate model of correlated noise (Adibi et al., 2014). In addition, light level-dependent changes in RGC signals and correlated noise may place important constraints on post-retinal computations across light levels. For example, downstream circuits that receive input from OFF-bt RGCs may fail to effectively process this input unless they too adapt their processing across light levels. Meanwhile, circuits that receive input from OFF-bs RGCs may be afforded a more static processing strategy. Thus, our results suggest that post-retinal areas may need to differentially process cell type inputs. Recent work elucidating LGN processing of RGC output confirm that some LGN neurons receive predominant input from a single type of RGC (Rompani et al., 2017; Liang et al., 2018; Roman Roson et al., 2019). These studies also find LGN neurons with input from diverse types of RGCs, posing further questions of how correlations between RGCs of different types may affect early visual processing.

Studies of light adaptation that span rod-to-cone signaling are relatively common in retina, but remain sparse in visual cortex. The few studies that have been performed suggest that V1 RFs are relatively invariant to changes in light level (Duffy and Hubel, 2007). The insights that RGC responses to the same stimulus change across light levels (Tikidji-Hamburyan et al., 2015), combined with the fact that correlated noise depends on light adaptation, motivates more research to understand the extent to which V1 and other regions can preserve an invariant representation of visual scenes across light levels.

## Acknowledgements

We would like to thank J. Pillow, J. Cafaro and S. Roy for helpful conversations, and L. Glickfeld, S. Lisberger and J. Mitchell for reading drafts of the manuscript. This research was supported by the A.P. Sloan Foundation (J.Z.), Canada Research Chairs program (J.Z.), Natural Science and Engineering Research Council of Canada (NSERC; J.Z.), the National Institutes of Health and National Eye Institute grants F31EY028833 (K.R.) and R01EY024567 (G.D.F), and the Whitehead Scholars Award (G.D.F).

## Contributions

K.R. and G.D.F. designed the study. Experiments and analysis were performed by K.R. with assistance from G.D.F. and J.Z. The manuscript was written by K.R. and G.D.F. with comments from J.Z.

## Methods

### MEA Recordings

All experiments were performed in accordance with the guidelines of Duke University’s Institutional Animal Care and Use Committee. Long-Evans rats were dark adapted overnight and euthanized with intraperitoneal injection of ketamine/xylazine followed by decapitation. Euthanasia and retinal dissections were performed in darkness with the assistance of infrared converters. We dissected dorsal pieces of retina that were approximately 3mm x 2mm large and placed them RGC-side down on an electrode array. The tissue was perfused with oxygenated Ames solution at a rate of 6-8 mL/min. Recordings were performed at 34°C. The MEA consisted of 512 electrodes with 60 μm spacing, covering an area of 0.9 x 1.8mm (Frechette et al., 2005; Ravi et al., 2018). The voltage on each electrode was sampled at 20 kHz and filtered between 80 and 2000 Hz.

### Visual Stimuli

Stimuli were presented with a gamma-corrected OLED display (SVGA+XL Rev3, Emagin, Santa Clara, CA). The image from the display was focused onto the photoreceptors using an inverted microscope (TiE, Nikon Instruments) with a 4x objective (CFI Super Fluor 4x, Nikon Instruments). Optimal focus was confirmed by presenting a high spatial resolution checkerboard noise stimulus (20×20 μm, refreshing at 15 Hz) and adjusting the focus to maximize the spike rate of RGCs over the MEA. The intensity of the stimulus was set using neutral density filters in the light path, and calibration was performed using previously described methods (Yao et al., 2018). In each recording, stimuli were first presented at the scotopic light level (1 Rh*/rod/s) while the retina was in a dark-adapted state. The tissue was adapted to the photopic light level (10,000 Rh*/rod/s) for 30 minutes before continuing recordings at that light level. The refresh rate of the stimulus was 60 Hz and 30 Hz at the photopic and scotopic light levels, respectively. The change in stimulus refresh offset effective contrast changes due to a ~2-fold increase in temporal integration of RGCs from the photopic to scotopic conditions. For GLM fitting and decoding, stimuli consisted of non-repeated, binary white noise interleaved with repeated, binary white noise segments (5 or 10s) to control for nonstationarities in recordings. Stimulus pixels in the checkerboard noise were squares with 240μm sides. For cell type classification, we followed previously described methods and presented drifting gratings and finer pixel checkerboard noise (60 μm squares, refreshing at 60 Hz) (Ravi et al., 2018).

### Spike Sorting and Neuron Identification

The spike sorting procedures have been described previously (Field et al., 2007). In brief, spikes on each electrode were identified by thresholding the voltage traces at 4 SD of a robust-estimate of the voltage SD. Spike sorting was performed by an automated PCA algorithm and verified by hand with a custom software (Shlens et al., 2006). Spike waveform clusters were identified as neurons only if they exhibited a refractory period (1.5ms) with <10% estimated contamination. To track identified RGCs across light conditions, cell clusters were sorted in the same PCA subspace at each light level. Neuron identity was verified by checking that electrical images (EIs) and RF locations were stable across conditions (Petrusca et al., 2007; Field et al., 2009). RGC types were classified at the photopic light level by first removing direction selective RGCs, and then clustering using RF properties and autocorrelation shapes (Ravi et al., 2018).

### Measuring noise correlations

Correlated noise was estimated by subtracting stimulus-driven correlations from the combined signal and noise correlations. Correlations were computed using responses to 100 (or 200) repeats of 10s (or 5s) white noise segments, binned at 5ms. First, the raw CCF was found by averaging the CCFs between two RGCs over all trials. Next, the shuffled CCF (shift predictor (Perkel et al., 1967)) was found by using spikes from one repeat for the first cell with spikes from a different repeat for the second cell. The shuffled CCF was averaged over all possible repeat combinations. Subtracting the shuffled CCF from the raw CCF gives the noise CCF. Correlation was quantified with the positive area under each correlogram, the full width of the peak, or the peak height at 0-time lag. The spatial scale of correlations for a population was found by fitting the data (e.g. Fig. 1E) to a single term exponential function. The coefficient of the exponential was the length scale of the correlations.

### Generalized Linear Model fitting

GLMs were fit separately at each light level. 50 minutes of non-repeated white noise were used to fit GLM parameters, with 100 (or 200) 10s (or 5s) segments of repeated white noise used for cross-validation. GLM RFs were approximated as rank one: they were composed of the outer product of a spatial filter and temporal filter (which approximated the spatial and temporal RFs, respectively). The temporal filter, spike history filter, and coupling filters were parameterized with a basis of 8 cosine functions. The nonlinearity used was the logexp2 function (https://github.com/pillowlab/GLMspiketools); no significant improvement was found using a spline nonlinearity. Only RGCs that had stable responses over the course of the recording were used in the GLM analysis (as judged by a consistent mean firing rate and uniform raster structure to repeated white noise sequences measured early and late in the experiment). For the coupled GLM fits, local populations of RGCs were chosen based on a central RGC and its neighbors. Only RGCs with an average firing rate above a threshold were included in the GLM; the threshold was given by the mean minus 1 SD of the firing rate of all recorded cells of a certain type. Across light levels, the same groups of RGCs were used to fit GLMs.

### Decoding

Following previous work (Pillow et al., 2008), GLM-based decoding was performed by computing the likelihood *p_j_* that a stimulus *x_j_* caused a recorded population response, where *x_j_* is one stimulus option of all possible binary white noise sequences. The decoded estimate comes from Bayes’ least squares estimate: 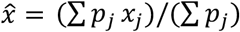. We decoded the intensity of one stimulus pixel over 6 frames in time, or 6 stimulus pixels in one frame for spatial decoding controls (Supp. Fig. 4). This decoding was repeated for 5000 trials on 16 minutes of non-repeated white noise data (held out from the fitting data). Decoding performance is reported with log SNR calculated from the mutual information between the decoded estimates and presented stimuli (Warland et al., 1997; Pillow et al., 2008). Bootstrapped SNRs for error bars were computed for each GLM using 500 subsamples of 3000 trials.

### Simple RF Model

The RF grid model consisted of 25-169 neurons with circular RFs arranged in a hexagonal grid. RF diameters were 250 μm, slightly larger than the stimulus pixel size of 240 μm. Cells responded linearly to white noise pixels according to the amount of overlap between that cell’s RF and the pixel. Each cell’s maximum firing rate was set to 30Hz. Noise correlation strength was set to decrease exponentially with distance between two cells and the coefficient of the exponential was the length scale of the correlations. We computed the discriminability of the center pixel’s intensity using *d*^2^ = Δ*μ^T^Q*^-1^Δ*μ*, where Δ*μ* is the vector of differences in mean neural firing rates between the decoded pixel value as black or white, and *Q* is the mean covariance matrix (Averbeck and Lee, 2006). To match the GLM decoding, here the covariance includes covariance of neurons and outer pixel intensities. This allowed conditioning the discrimination on the intensities of non-decoded pixels. Thus, decoding was performed given the population responses and intensity of outer stimulus pixels. The coupled decoder utilized the full covariance matrix, while the independent decoder ignored non-diagonal entries in the neuron-neuron covariance block.

### Statistics

Significance tests were performed using Student’s paired sample t test (for comparisons of one cell type across light levels) and Student’s 2 sample t test (for comparisons across cell types).

## Supplementary Figures

**Supp. Figure 1:**
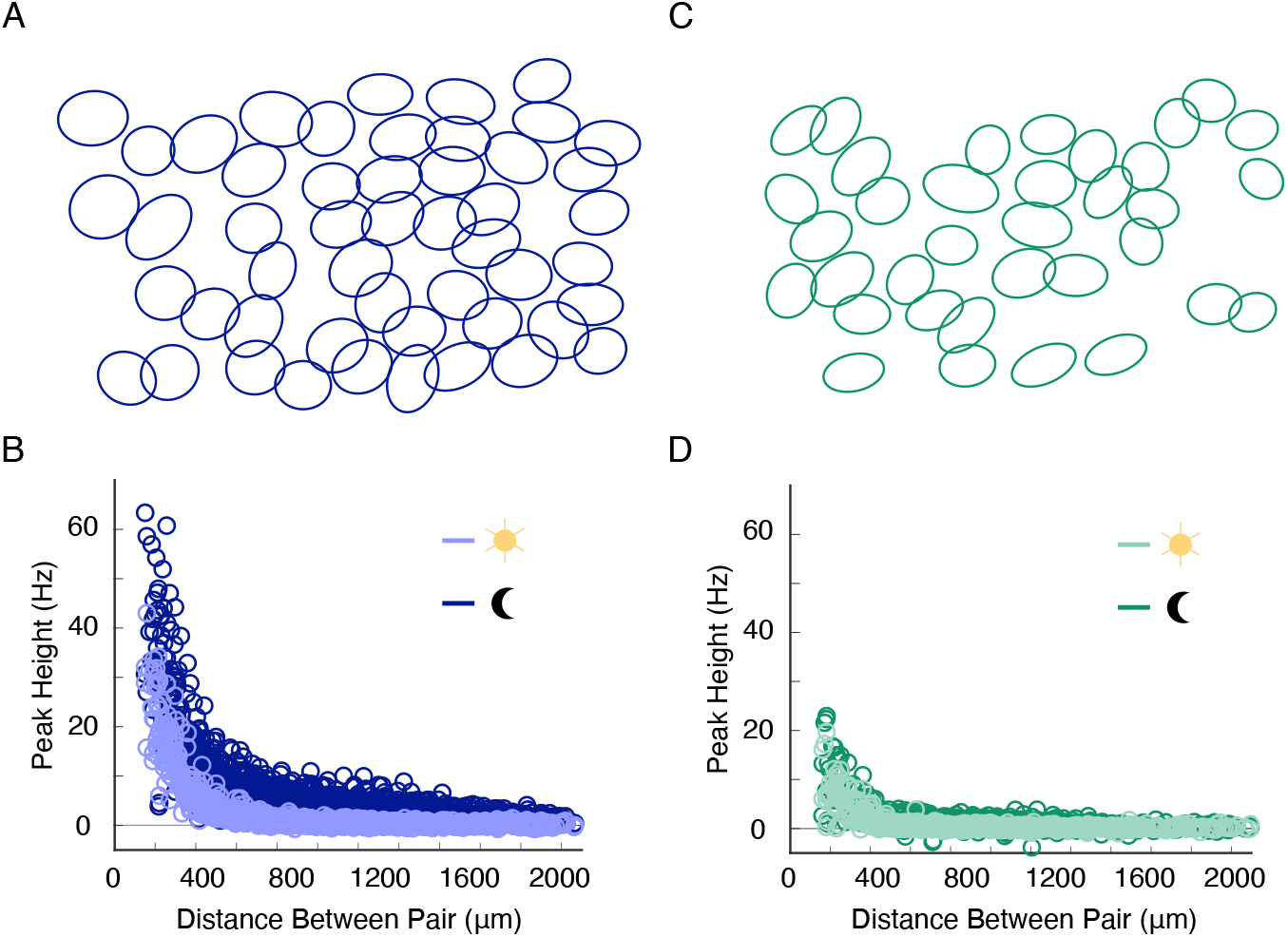
Spontaneous noise correlations depend on light level and cell type. **A.** RF mosaic of OFF-bt RGCs. **B.** Strength of noise correlations, measured with a static, full-screen black stimulus, over pairwise distances for the OFF-bt RGC population. Each point shows the cross-correlogram height at 0-time lag for a given pair of cells (1225 RGC pairs from 1 retina; photopic light level: 1000 Rh*/rod/s; scotopic light level: 0.1 Rh*/rod/s). **C & D**. same as **A** & **B** for OFF-bs RGCs (595 RGC pairs from 1 retina).

**Supp. Figure 2:**
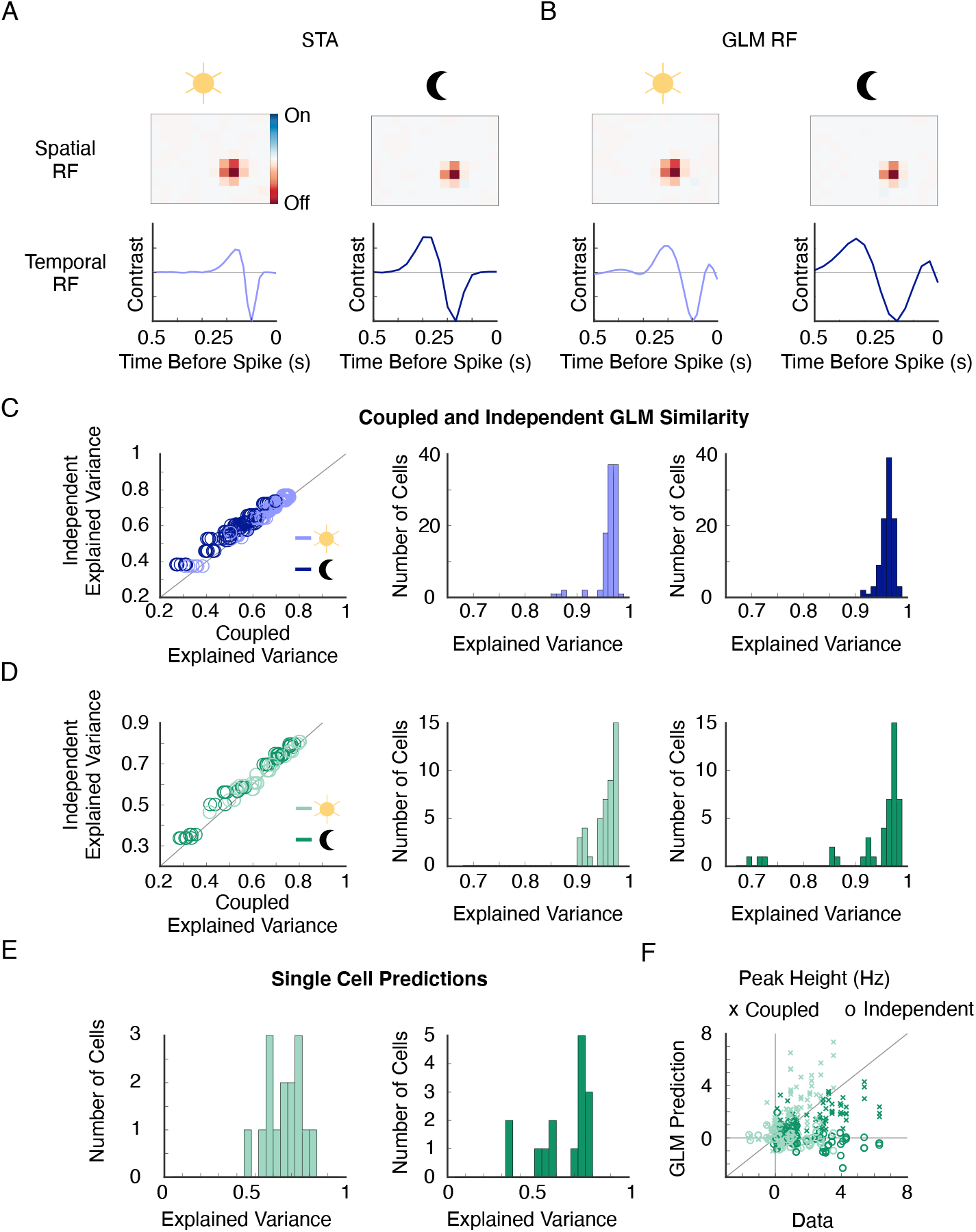
Summary of GLM fitting performance. **A.** Spike triggered average (STA) of an example OFF-bt RGC across light levels. Note that these RGCs have space-time separable RFs (Ravi et al., 2018). Top row, spatial components of the STA estimate the spatial RF. Bottom row, time courses of the STA estimate the temporal RFs (also called temporal filters). Notice the slower time course at the scotopic light level. **B.** Same as **A** but spatial and temporal filters are from the GLM fit. **C.** Comparing coupled and independent GLM performances for OFF-bt RGCs. Left, performances in predicting firing rates are similar for the coupled and independent GLMs (101 cells from 16 groups of RGCs from 1 retina; all data: photopic: independent GLM explained variance = 0.59 ± 0.013, mean *±* s.e.m., 97 RGCs from 4 retinas, coupled GLM explained variance = 0.59 *±* 0.007, 55 groups of RGCs from 4 retinas, scotopic: independent GLM explained variance = 0.58 ± 0.01, 66 RGCs from 3 retinas, coupled GLM explained variance = 0.51 *±* 0.008, 37 groups of RGCs from 4 retinas). Most RGCs were used once in the independent GLMs but are part of multiple coupled GLMs. Middle, distribution of explained variance between coupled and independent PSTH predictions at the photopic light level (all data: 0.94 ± 0.002). Right, distribution of explained variance between coupled and independent PSTH predictions at the scotopic light level (all data: 0.94 ± 0.002). **D.** Same as **C** for OFF-bs RGCs (44 cells from 8 groups of RGCs from 1 retina; all data: photopic: independent GLM explained variance = 0.51 ± 0.02, 69 RGCs from 4 retinas, coupled GLM explained variance = 0.52 *±* 0.01, explained variance between independent and coupled GLMs = 0.9 *±* 0.006, 37 groups of RGCs from 4 retinas, scotopic: independent GLM explained variance = 0.65 ± 0.02, 42 RGCs from 3 retinas, coupled GLM explained variance = 0.63 *±* 0.01, explained variance between independent and coupled GLMs = 0.93 *±* 0.005, 20 groups of RGCs from 3 retinas). **E.** Distribution of explained variances for the GLM predicted PSTHs in OFF-bs RGCs (15 RGCs from 1 retina). **F.** Cross-correlogram peak predictions for the independent and coupled GLMs across the OFF-bs RGC population (102 pairs from 9 groups of RGCs from 1 retina).

**Supp. Figure 3:**
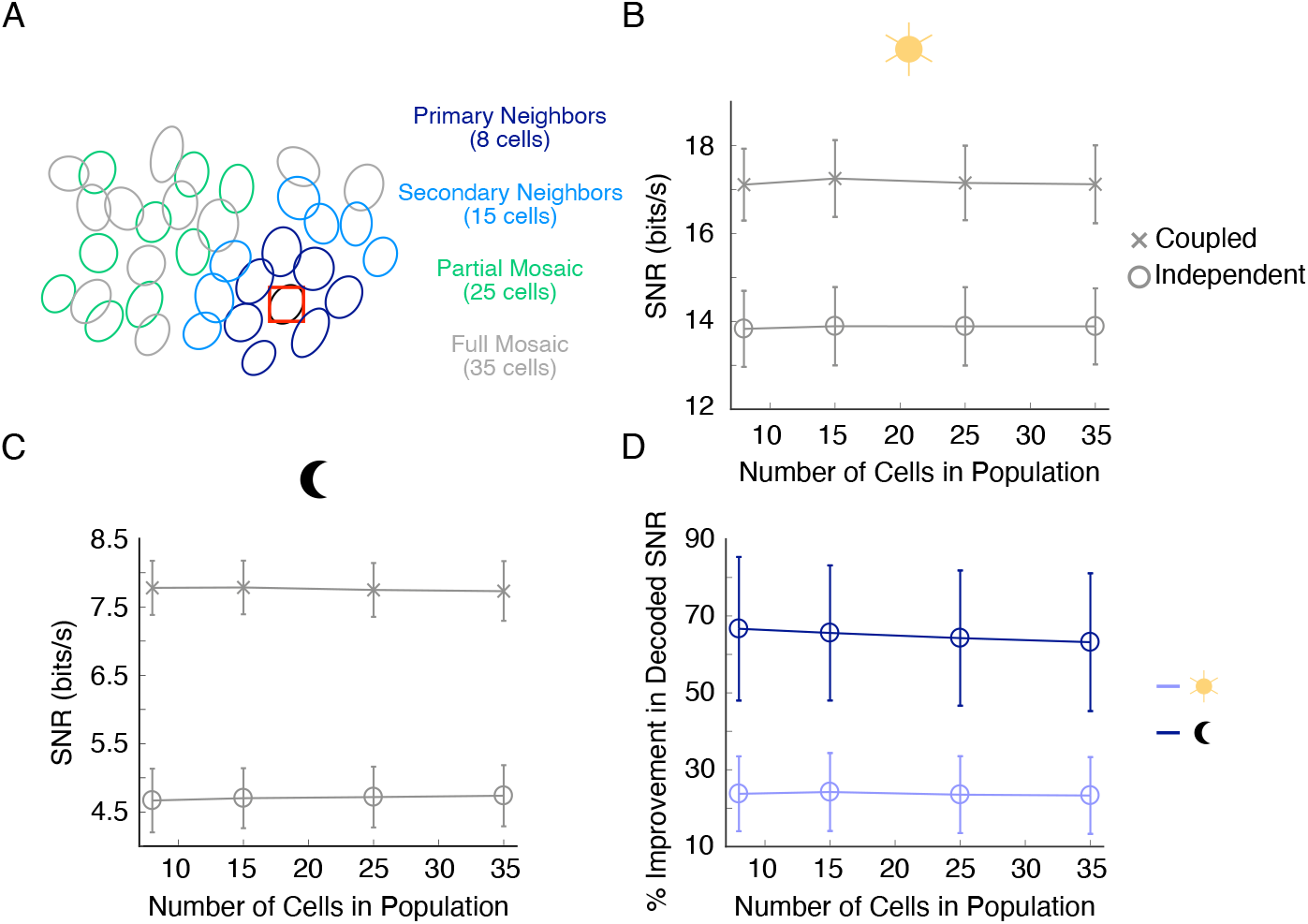
Including RGCs beyond immediate neighbors does not significantly impact temporal decoding results. **A.** OFF-bt RF mosaics colored to show the four different populations that were compared with GLM decoding. The first group included an RGC centered over the decoded stimulus pixel and all of its primary neighbors. The second group adds secondary neighbors, the third group adds some far away RGCs, and the fourth group uses the whole recorded population. **B.** Decoded SNR for the independent and coupled GLMs fit with the groups in **A** at the photopic light level. Error bars (SD) come from bootstrapping SNR. **C.** Same as **B** for the scotopic light level. **D.** Percent improvement in decoded SNR between the coupled and independent GLMs for the different RGC groups.

**Supp. Figure 4:**
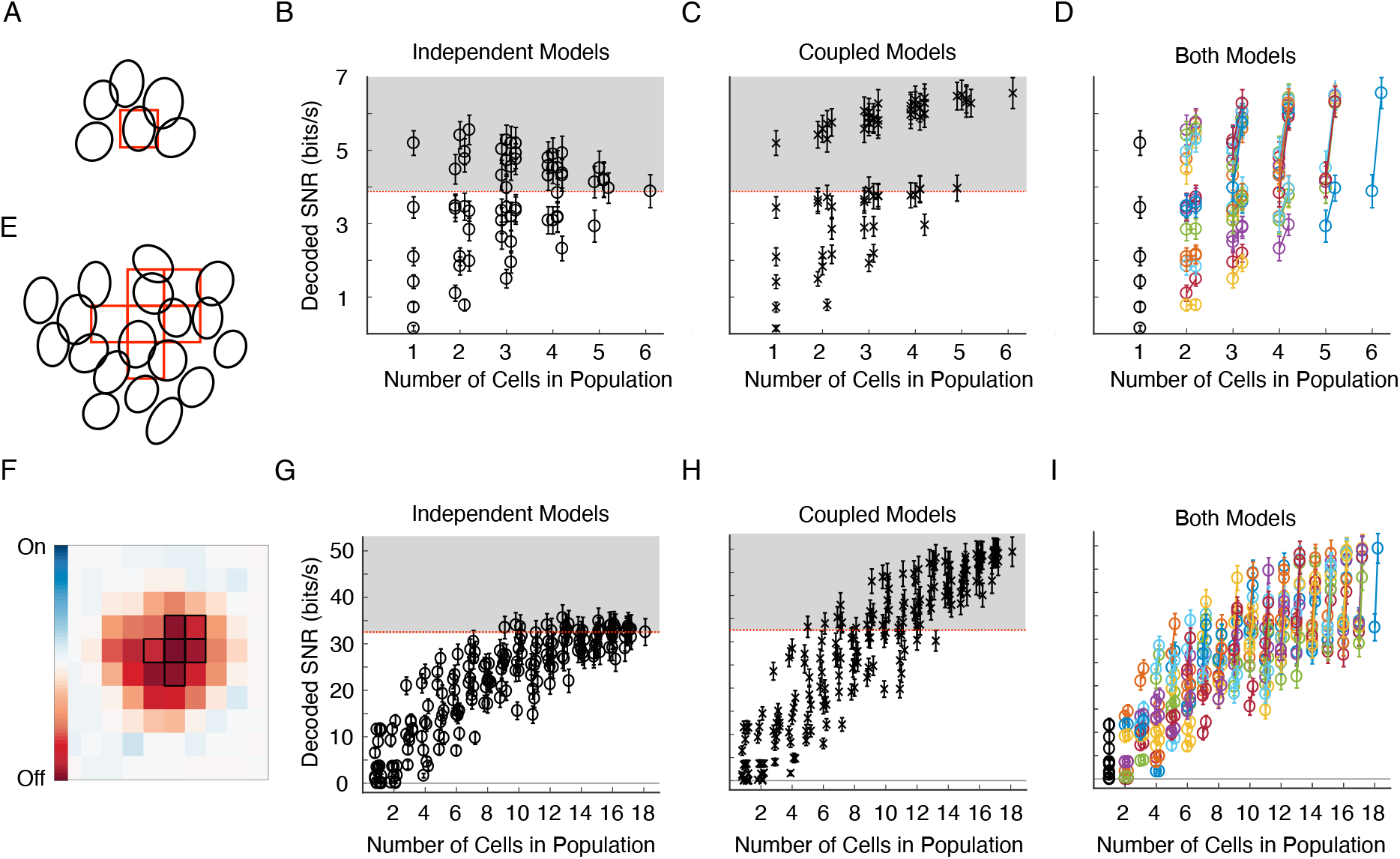
Population failure as a function of group size and decoding in time (panels A-D) or space (panels E-I). **A.** RF mosaic for 6 OFF-bt RGCs used to decode the temporal sequence of the stimulus pixel in red. **B.** Decoded SNR for independent GLMs as a function of number of cells included in the population. All possible combinations of 1-6 RGCs were used. The independent GLM with all 6 cells decodes less information than GLMs with some combinations of 1-6 RGCs. Points are slightly jittered in the x direction for visualization. **C.** Same as **B** for coupled GLMs. Several combinations of 2-6 coupled GLMs (including 1 single cell GLM) decode more information than the independent GLM with 6 cells (gray region). **D.** Decoded SNR for independent and coupled GLMs, where models using the same group of RGCs are connected by a line. For visualization, the coupled SNRs are plotted slightly to the right of the independent SNRs. **E.** RF mosaic for 18 OFF-bt RGCs used to decode the spatial pattern of stimulus pixels in red. Here we consider when an independent GLM consisting of many RGCs decodes worse than a GLM made up of a smaller population. This scenario exhibits population failure because the decoder with more cells fails to take advantage of the information provided by larger population input. Expanding to larger populations in this way is necessary because of the large size of stimulus pixels relative the RGC RFs, which makes it unlikely that a single RGC can decode a large spatial pattern well. **F.** Cumulative RF coverage for the 18 RGCs. The spatial pattern of stimulus pixels that was decoded is outlined in black. This plot shows that the decoded stimulus pixels are well represented by the group of RGCs. **G.** Decoded SNR for independent GLMs as a function of number of cells included in the population. For groups with 2-17 RGCs, all possible combinations of RGCs were subsampled. Note that decoded SNR appears to plateau at ~15 cells. **H.** Same as **G** for coupled GLMs. The independent GLM using all 18 RGCs decodes less information than coupled GLMs with some combinations of 7, 9-17 RGCs (gray region). Note that decoded SNR would likely continue growing with a larger population of RGCs. **I.** Decoded SNR for independent and coupled GLMs, where models using the same group of RGCs are connected by a line. For visualization, the coupled SNRs are plotted slightly to the right of the independent SNRs. For a population of 18 RGCs, the coupled model performs 52 ± 16.61 % (mean ± SD) better than the independent model.

**Supp. Figure 5:**
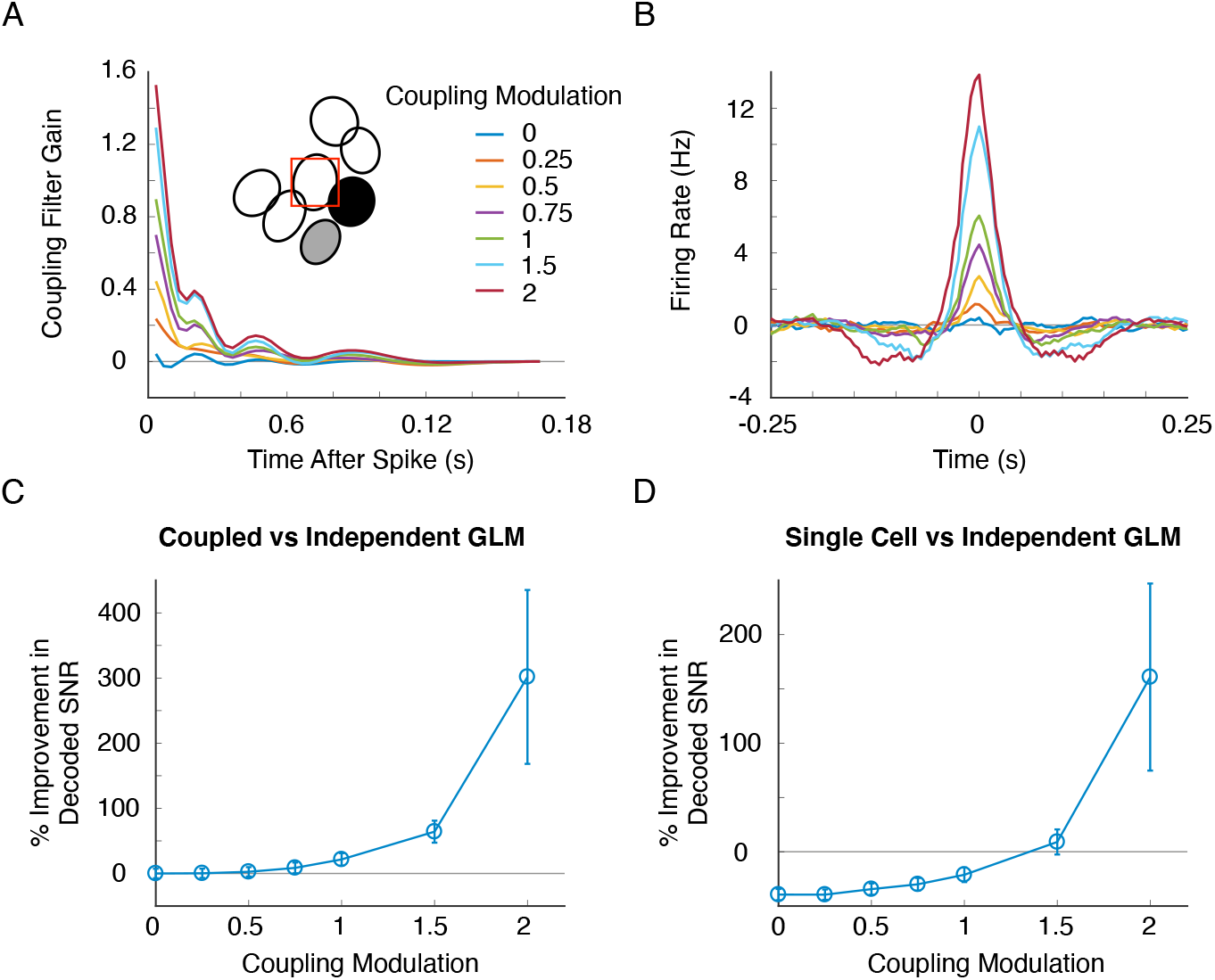
Stronger coupling causes a greater improvement in decoding when accounting for correlations relative to assuming independence. We started with a coupled GLM fit to a population of OFF-bt RGCs at the scotopic light level (inset in **A**). We then altered all coupling filters in the model by a constant factor, simulated spike trains, refit coupled and independent GLMs and decoded stimuli from the simulated spike trains to determine the impact of directly altering coupling strengths on decoding performance. **A.** Coupling filters between two RGCs in the population (highlighted in the inset) for the indicated coupling modulation factors. These coupling filters were used in the GLM to simulate responses. **B.** Resulting noise CCFs from simulated spike trains of the gray and black cells in A). **C.** Percent improvement in decoded SNR between the coupled and independent GLM as a function of coupling modulation. Stronger coupling filters make ignoring correlated activity more deleterious for decoding. **D.** Same as but comparing decoding with a single cell and the independent GLM to demonstrate population failure. Population failure does not occur for the original group of cells (coupling modulation = 1), but increasing coupling strength causes the independent GLM to decode less information than the single cell.

**Supp. Figure 6:**
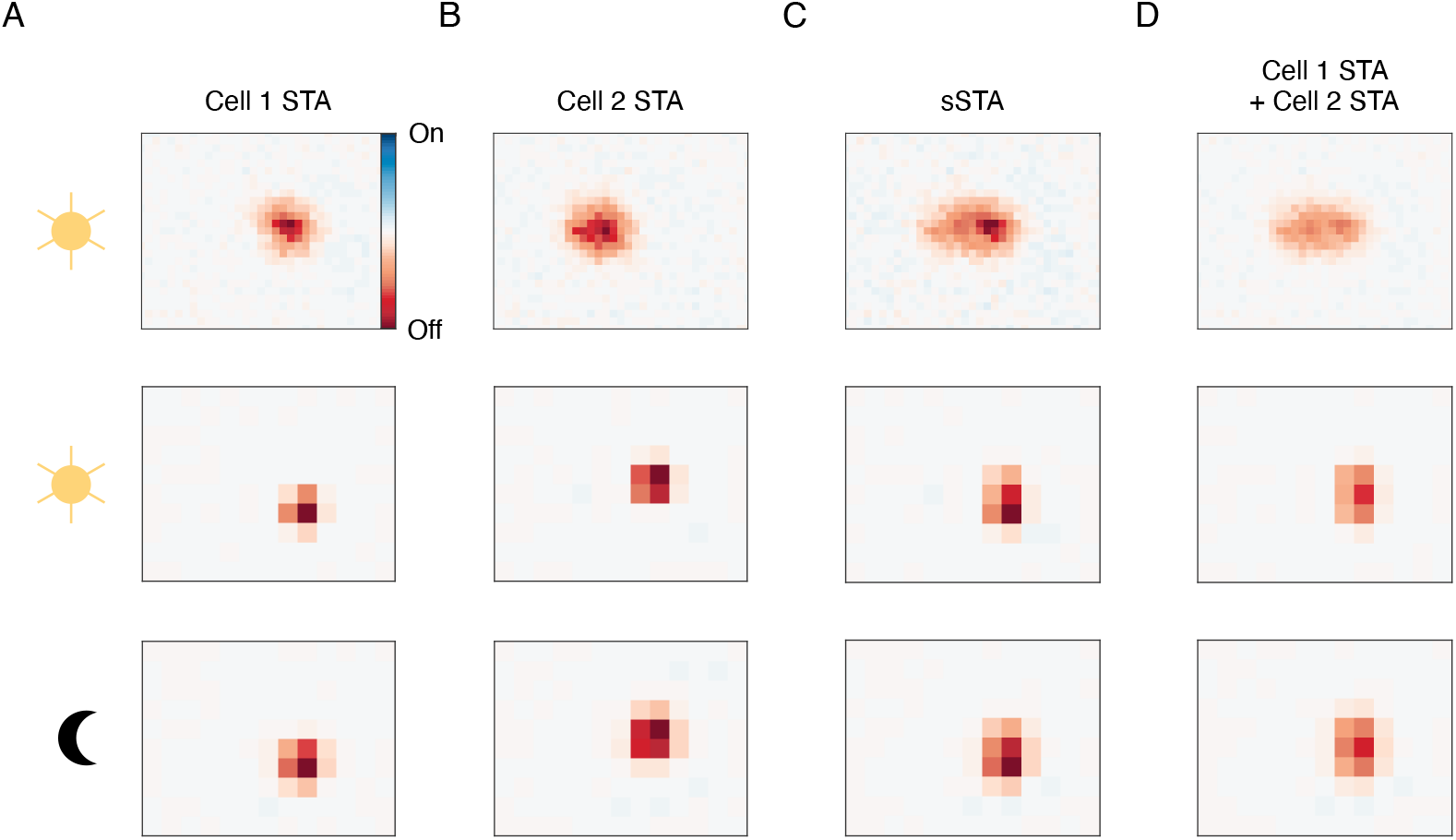
Synchronous spikes between pairs of RGCs do not encode finer spatial information. A. Top, spatial RF of an example OFF-bt RGC at the photopic light level measured with a fine spatial resolution. Middle, the same cell’s RF at the spatial resolution used for GLM decoding at the photopic light level. Bottom, the cell’s RF at the scotopic light level. B. RFs for an RGC neighboring the cell in A. C. sSTAs between the RGCs of A & B. The sSTAs do not resemble the intersection of the two individual RFs. D. The union of the RFs from A & B.

